# Shared effect modeling reveals that a fraction of autoimmune disease associations are consistent with eQTLs in three immune cell types

**DOI:** 10.1101/053165

**Authors:** Sung Chun, Alexandra Casparino, Nikolaos A Patsopoulos, Damien Croteau-Chonka, Benjamin A Raby, Philip L De Jager, Shamil R Sunyaev, Chris Cotsapas

**Author notes:** correspondence to SRS and.

## Abstract

The majority of autoimmune disease risk effects identified by genome-wide association studies (GWAS) localize to open chromatin with gene regulatory activity. GWAS loci are also enriched for expression quantitative trait loci (eQTLs), suggesting that most disease risk variants exert their pathological effects by altering gene expression^1,2^. However, because causal variants are difficult to identify and *cis*-eQTLs occur frequently, it remains challenging to translate this bulk observation into specific instances of a disease risk variant driving changes to gene regulation. Here, we use a novel joint likelihood framework with higher resolution than previous methods to identify loci where disease risk and an eQTL are driven by a single, shared genetic effect as opposed to distinct effects in close proximity. We find that approximately 25% of autoimmune disease loci harbor an eQTL driven by the same genetic effect, but the majority of loci do not. Thus, we uncover a fraction of gene regulatory changes as strong mechanistic hypotheses for disease risk, but conclude that most risk mechanisms do not involve changes to basal gene expression.

The autoimmune and inflammatory diseases (AID) – multiple sclerosis, type 1 diabetes, inflammatory bowel disease, and more than 80 others – are highly heritable, complex diseases that cumulatively afflict 8% of the population^3,4^. Pathology is driven by loss of tolerance to self-antigens, resulting in either systemic or tissue-specific immune attack. They are extensively co-morbid^5,6^ with extensive sharing of AID genetic risk^7–9^, indicating that pathogenic mechanisms are also shared^10^. Disease mechanisms are still poorly understood, so current therapies control symptoms rather than root causes, often by suppression of major immune responses. Consortium-driven genetic mapping studies have identified hundreds of genomic regions mediating risk to several AID, and these communities have collaborated to develop the ImmunoChip custom genotyping array to deeply interrogate 185 of these loci^11^.

These disease associations are primarily non-coding: lead GWAS SNPs are more likely to be associated with expression levels of neighboring genes than expected by chance^12^, and the same lead SNPs are enriched in regulatory regions marked by chromatin accessibility and modification^1,13^. Fine-mapping reveals enrichment of AID-associated variants in enhancer elements specifically active in stimulated T cell subpopulations^14^, and formal disease heritability partitioning analyses also show strong enrichment in such regions of gene regulatory potential^15,16^. Collectively, these strands of evidence suggest that the majority of disease risk is mediated by changes to gene regulation in specific cell subpopulations. However, these bulk overlap and enrichment analyses do not formally assess whether expression levels and disease risk are associated to the same underlying variant or are due to independent effects in the same region^17,18^. Several methods have been developed to identify pathogenic genes within GWAS loci relying on eQTL co-localization^19–22^. However, these are limited in resolution to detect situations where causal variants for the disease trait and the eQTL are distinct but in linkage disequilibrium.

Here, we present an approach to test if a GWAS risk association and an eQTL are driven by the same underlying genetic effect, accounting for the LD between causal variants. Using data from ImmunoChip studies of seven AID comprising >180,000 samples in total (Table S2), we apply this approach to test if associations in 272 known risk loci are consistent with *cis*-eQTL for genes in each region, measured in three relevant immune cell populations: lymphoblastoid cell lines (LCLs), CD4^+^ T cells and CD14^+^ monocytes^23,24^.

When associations to two traits in a locus – here, a disease trait and an eQTL – are driven by the same underlying causal variant, the joint evidence of association should be maximized at the markers in tightest LD with the (potentially unobserved) causal variant^18,25^. Here, we directly evaluate the joint likelihood that both trait associations are due to the same underlying causal variant (Figure S1), unlike previous approaches that look for similarities in the shape of the association curve over multiple markers^19,20,26,27^. We expect that when the underlying causal effect is shared, joint likelihood is maximized when we model the same causal variant in both traits; conversely, when the underlying causal variants are different, we expect maximum joint likelihood when we model their closest proxies. We empirically derive the null distribution of the joint likelihood ratio statistic by comparing disease associations to permuted eQTL data. The resulting *P* value is asymptotically conservative against the null of distinct causal variants, as the likelihoods of two competing models will be further contrasted with increasing sample and effect sizes (Supplementary Notes). We are thus able to directly evaluate whether associations in the same genomic locus for two traits, observed in different cohorts of individuals, are due to the same underlying causal variant.

To assess the performance of our method, we benchmarked it against *coloc*^19^, a well-calibrated Bayesian framework that considers spatial similarities in association data across windows of markers in a locus. We simulated pairs of case-control cohorts with either the same or distinct causal variants driving association in each. Though both methods had excellent performance in simulations where the two distinct causal variants are in complete linkage equilibrium (AUC=0.99 for both when ***r^2^*** < 0.5), compared to *coloc* we found that our method maintained higher specificity even as the linkage disequilibrium between distinct causal variants became high (AUC = 0.92 compared to 0.70 for *coloc* when 0.7 < ***r^2^*** < 0.8; Figure S3, Tables S3 and S4). In practice, our resolution becomes limited at very high LD levels (***r^2^***>0.8), and we are unable to reliably distinguish between two causal variants in very high LD and a single causal variant associated to both traits. Thus, within these limits, we can accurately detect cases of shared genetic effects between two traits.

We first identified densely genotyped ImmunoChip loci showing strong association to each disease from publicly available summary data (immunobase.org; Table 1). These include associations to both Crohn disease and ulcerative colitis, collectively designated inflammatory bowel disease, and to each disease alone^28^. Due to the extensive LD and complex natural selection present in the Major Histocompatibility Locus, we excluded this region from consideration. We next identified genes in a 1Mb window centered on the most associated variant in each locus. Consistent with previous observations that eQTLs are frequently found in GWAS loci, we found that all loci but 11 had at least one gene with an eQTL (p < 0.05) in at least one of the three cell types, with most such effects common across all three tissues (Table 1). In total, we found 9,268 pairs of disease and eQTL associations across 261 ImmunoChip loci. We then tested each of these pairs with our joint association likelihood method to assess if the eQTLs appear driven by the same underlying effect as the disease associations. We find evidence for shared effects for only 57/9,268 pairs in 41/261 loci across all diseases, with the proportion varying from 2/40 (5%) for type 1 diabetes loci to 4/10 (40%) for ulcerative colitis loci (false discovery rate < 5%; Tables 1 and 2). Of these 57 shared effects, 43 pass even the more stringent family-wise multiple testing correction (Bonferroni corrected P < 0.05). Thus, our analysis reveals that in the majority of AID loci, variants causally involved in disease phenotypes do not overlap variants responsible for eQTL signals. Overall, we find that only a small minority of tested disease-eQTL pairs in the same locus show any evidence of a shared association, whereas >75% show evidence of being driven by distinct genetic variants in the same locus (Figure 1).

**Table 1.** Only a minority of disease associations share causal variants with eQTLs across three immune cell subpopulations. We identified 261 disease associations in ImmunoChip regions with at least one eQTL within 100kb of the most associated SNP. Only 41/261 (16%) of these associations show evidence of a shared effect with an eQTL in that region. Thus, while eQTLs are abundant in disease-associated loci, they do not appear to be driven by the same causal variant as the disease association. ^a^We only consider associations reported at genome-wide significant levels and overlapping genomic regions densely genotyped on ImmunoChip, excluding conditional peaks and MHC loci (see Methods). ^b^eQTLs are selected if there is nominal association (eQTL p < 0.05) to at least one SNP within 100kb of the most associated SNP to disease, and a transcription start site of the gene within 1Mb of that SNP. cNumber of loci where disease association is consistent with a shared effect for at least one eQTL (FDR < 5%).

| Disease | Number of loci |  |  |  |  |  |  |  |  |
| --- | --- | --- | --- | --- | --- | --- | --- | --- | --- |
|  | Densely<br>genotyped <sup>a</sup> | eQTL present <sup>b</sup> |  |  |  | Driven by same effect <sup>c</sup> |  |  |  |
|  |  | CD4<sup>+</sup> | CD14<sup>+</sup> | LCL | Total | CD4<sup>+</sup> | CD14<sup>+</sup> | LCL | Total |
| **MS** | 59 | 54 | 55 | 55 | 56 | 8 | 3 | 6 | 12 |
| **IBD** | 69 | 69 | 69 | 68 | 69 | 6 | 9 | 1 | 12 |
| **Crohn** | 19 | 18 | 18 | 18 | 18 | 2 | 1 | 0 | 3 |
| **UC** | 10 | 10 | 9 | 10 | 10 | 2 | 1 | 3 | 4 |
| **T1D** | 47 | 39 | 40 | 36 | 40 | 2 | 0 | 0 | 2 |
| **RA** | 34 | 34 | 34 | 34 | 34 | 2 | 0 | 1 | 3 |
| **CEL** | 34 | 34 | 34 | 34 | 34 | 3 | 2 | 0 | 5 |
| **Overall** | 272 | 258 | 259 | 255 | 261 | 25 | 16 | 11 | 41 |

**Table 2.**
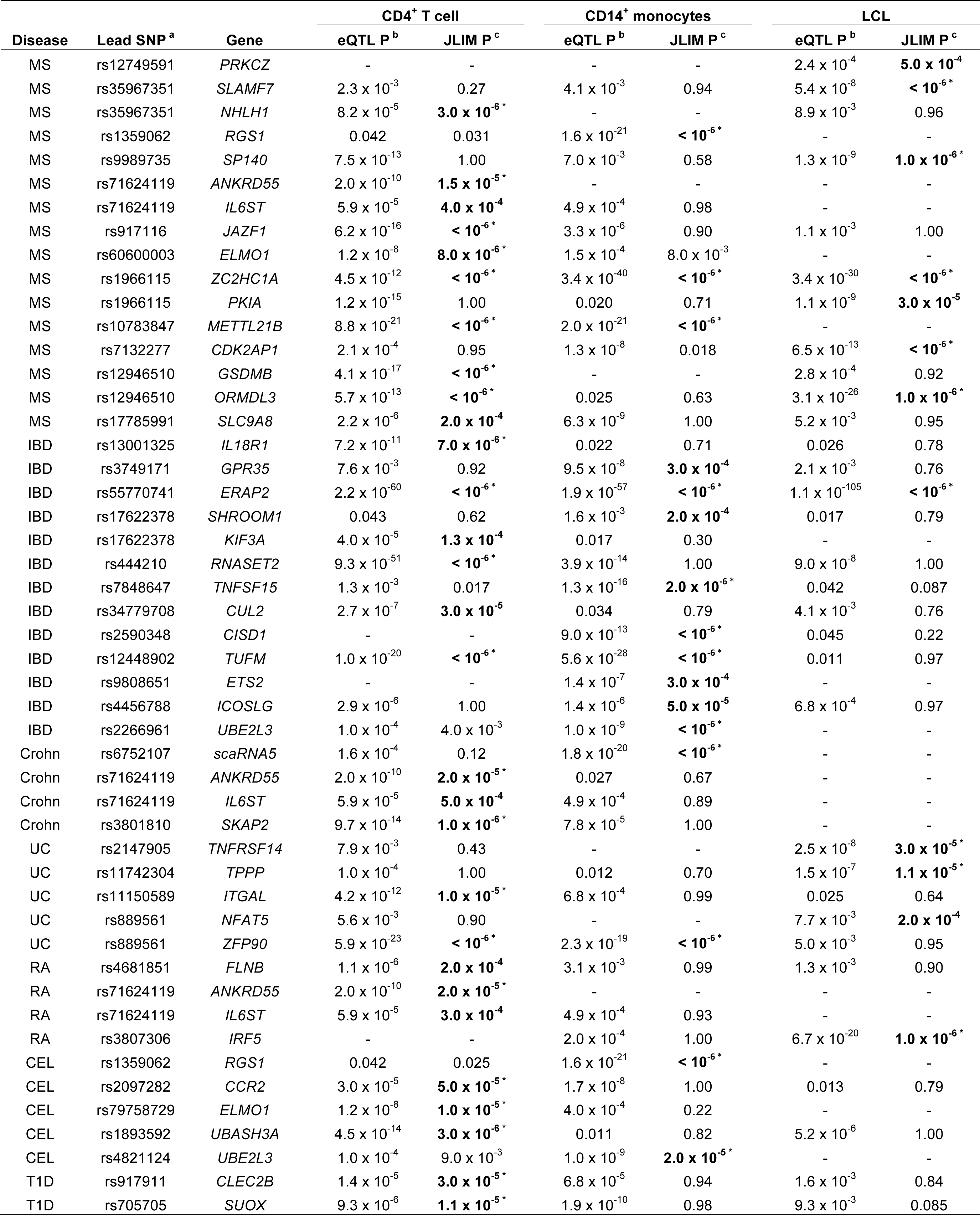
Forty one loci harbor eQTLs driven by the same variants as an association to at least one of seven diseases. We find 57 instances of shared disease-eQTL effects in 41 loci (joint likelihood of shared association FDR < 5%). ^a^Variant with the minimum association *p* value to disease in the ImmunoChip summary statistics. ^b^Minimum eQTL *p* value for any SNP within 100kb of the lead SNP. Dashes (-) indicate genes that are either not detected or with minimum eQTL P > 0.05 in that cell type. ^c^Highlighted in bold are disease-eQTL pairs with false discovery rate < 5%. Asterisk (*) marks eQTL genes passing Bonferroni correction.

**Figure 1.**
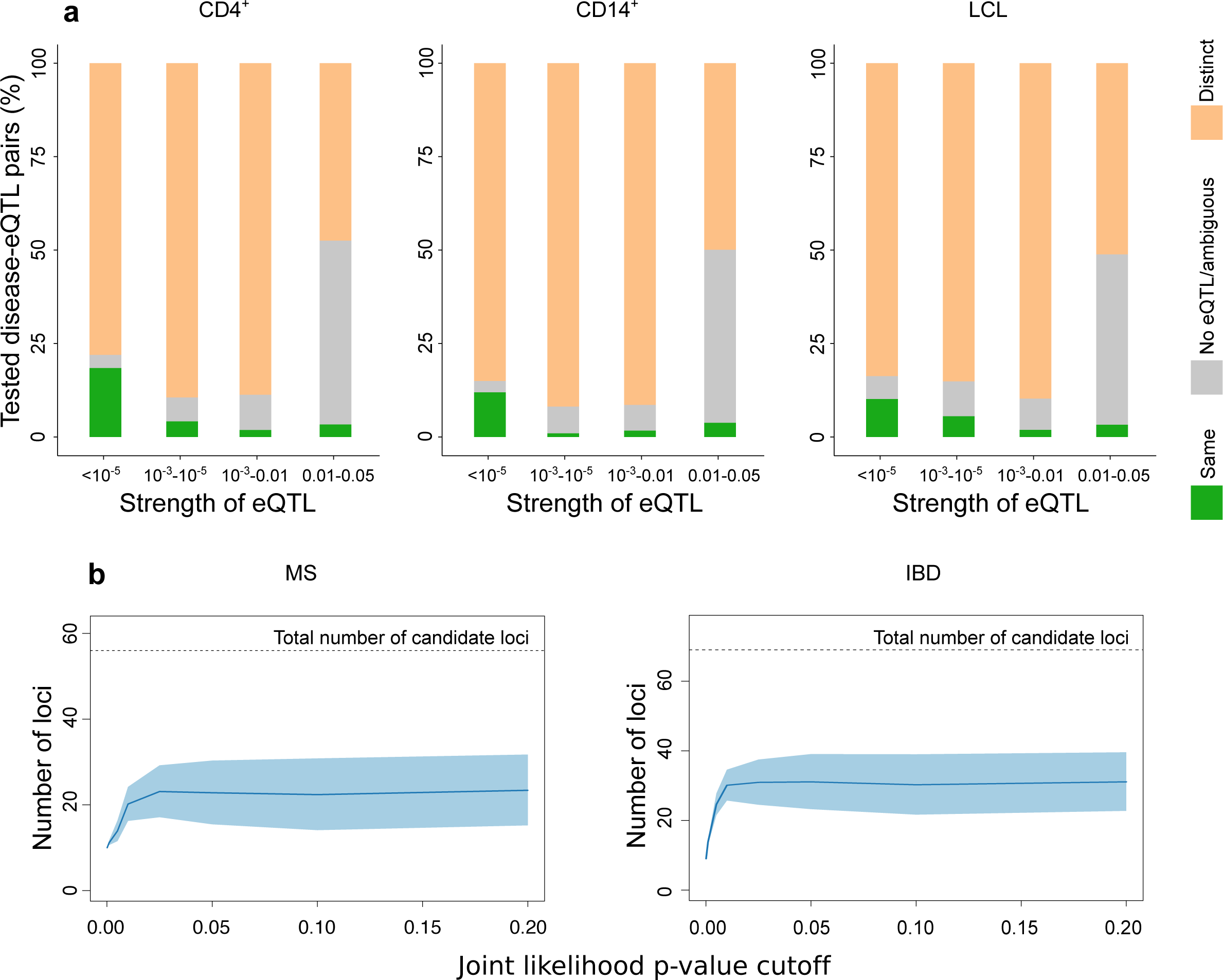
Only a minority of disease associations share causal variants with eQTLs across three immune cell subpopulations. **(a)** We find strong evidence that approximately 75% of eQTLs are driven by distinct causal variants (orange) to 261 disease risk associations across 155 ImmunoChip regions. The strength of eQTL association does not influence the proportion of shared effects (green) we are able to detect, suggesting this lack of overlap is not due to lack of power. We find no compelling evidence for either shared or distinct associations for a small proportion of disease-eQTL pairs (gray). **(b)** The median number of loci with at least one shared effect eQTL in any tissue (blue line) at more liberal significance thresholds remains constant after false positive adjustment, further supporting this conclusion. The shaded area represents the lower and upper expectation bounds for disease-eQTL pairs driven by the same causal variant. Only 31-57% of multiple sclerosis associations and 37-57% of inflammatory bowel disease associations are consistent with eQTL effects. Equivalent data for the other diseases are presented in Figure S14.

We sought to explain this lack of overlap between disease associations and eQTLs, despite their frequent co-occurrence in the same loci. In particular, although our method showed good performance in simulated data (Figure S4), we remained concerned that this lack of overlap may be due to low statistical power in the eQTL data, which come from cohorts of limited sample size. However, we find that even amongst the most strongly supported eQTLs (p < 10^−5^), < 25% show evidence of shared effects with disease associations. Conversely, we find strong evidence for distinct effects for the majority of disease-eQTL pairs, with only a subset of comparisons being ambiguous, suggesting that our method is adequately powered to detect shared effects where they exist (Figures 1a and S11-13). To assess whether power affects the total number of loci, rather than eQTL, that can be resolved, we looked more deeply at our significance threshold settings. We find that more liberal thresholds do not increase the number of true positive results after adjusting for false positive rate, indicating that most loci do not contain *any* gene with an eQTL consistent with the disease association (Figures 1b and S14). Cumulatively, our results demonstrate that only a minority of AID risk effects drive eQTLs in the three cell populations we tested, which are drawn from diverse lineages of the immune system.

We next focused on the subset of 57 disease/eQTL pairs in 41 loci where we could detect strong evidence of a shared effect (Table 2). We find that 51/57 (89%) of effects are restricted to one cell population, indicating that tissue-specific eQTLs are important components of the molecular underpinnings of disease (Figures S5 and S6). The remaining six effects are detected in multiple cell populations; for example, the multiple sclerosis association at rs10783847 on chromosome 12 is consistent with eQTLs for the transcript of methyltransferase-like 21B (*METTL21B*) in both CD4^+^ T cells and CD14^+^ monocytes, but not with eQTLs for the remaining 38 genes in the immediate locus (Figure 2). Although *METTL21B* is expressed in LCLs, there is no evidence of an eQTL in this tissue within 1Mb from rs10783847. Similarly, for the multiple sclerosis association at rs1966115 on chromosome 8 and eQTLs for *ZC2HC1A*, and for the inflammatory bowel disease association at rs55770741 on chromosome 5 and eQTLs for *ERAP2*, we detect a shared effect in all three cell populations. In several cases we find tissue-specific shared effects despite strong eQTLs for the same gene in other tissues: for *TUFM* and inflammatory bowel disease risk at rs12448902 on chromosome 16, we find shared effects in CD4^+^ and CD14^+^ but not LCLs, where we see a *TUFM* eQTL at p = 0.01 (joint likelihood P = 0.97). For *ZFP90* and ulcerative colitis risk at rs889561 on chromosome 16, we also find shared effects in CD4^+^ and CD14^+^ but not LCLs, where we observe a *ZFP90* eQTL at p = 0.005 that has a low likelihood of shared effect with GWAS (joint likelihood P = 0.95). Instead, we find evidence of sharing between disease risk and an eQTL for *NFAT5* in LCLs. Thus, despite the presence of eQTLs for a gene in multiple tissues, not all these effects are consistent with disease associations suggesting that disease-relevant eQTLs are tissue specific.

**Figure 2.**
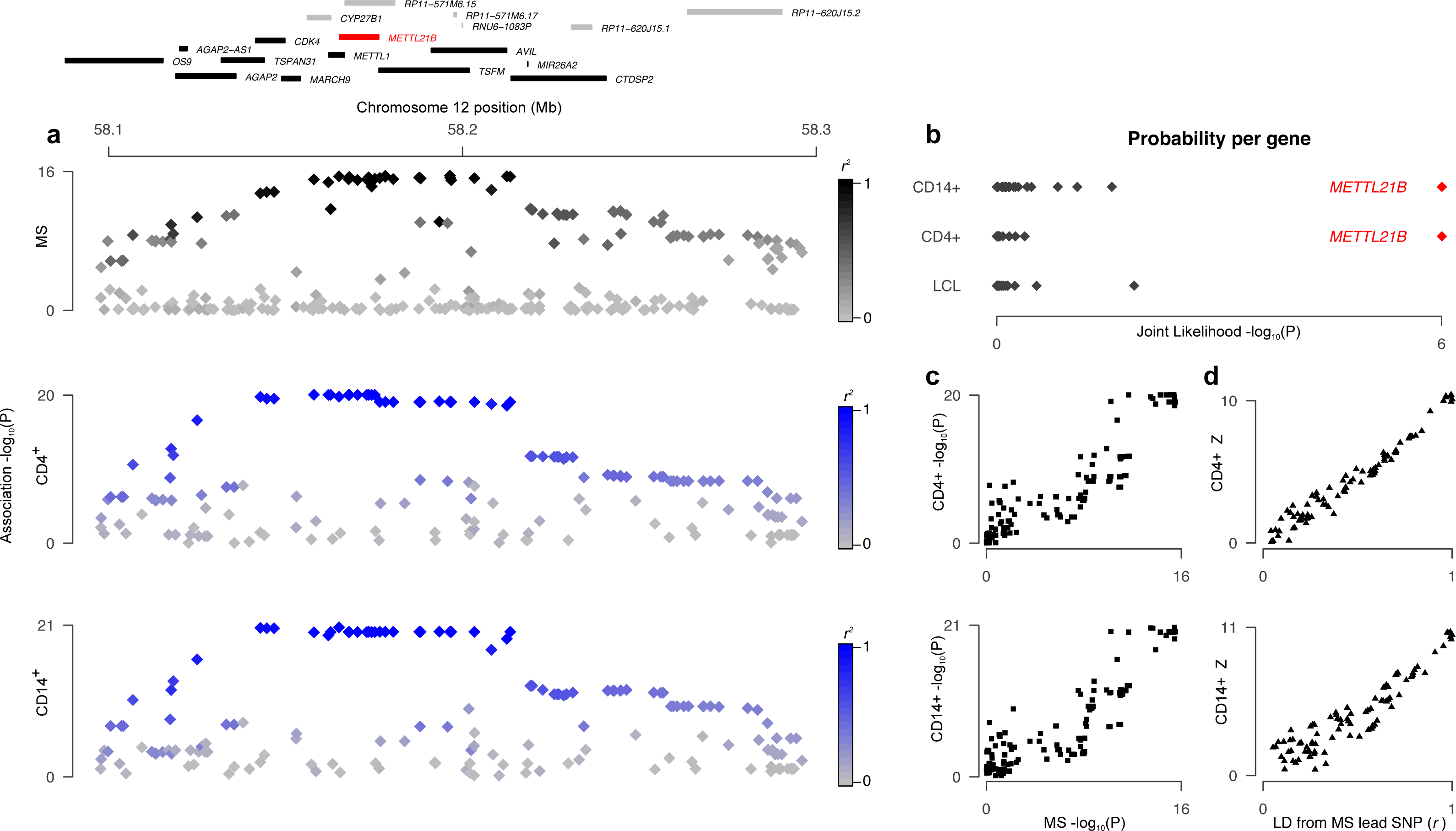
A multiple sclerosis association on chromosome 12 is consistent with eQTLs for *METTL21B* in both CD4^+^ T cells and CD14^+^ monocytes. **(a)** A genome-wide significant association to multiple sclerosis risk (upper panel; shading denotes strength of LD to the most associated variant rs10783847). This association is consistent with eQTLs for *METTL21B* in CD4^+^ T cells (middle panel) and CD14^+^ monocytes (lower panel, both shaded by LD to rs10783847), but not to eQTL data for any other genes in the region (upper gene track: black boxes denote 38 genes with eQTL data available in addition to *METTL21B* (red); gray denotes genes which are not reliably detected in our data or do not have eQTL p < 0.05 in the region). **(b)** Joint likelihood p-values for 39 candidate genes analyzed for this MS association peak in three cell types. Those with FDR < 5% are shown in red. **(c)** Association p-values for MS risk (x-axis) and eQTLs (y-axis) are strongly correlated for both CD4^+^ T cells (middle panel) and CD14^+^ monocytes (lower panel). **(d)** Similarly, eQTL association Z statistics scale linearly with LD (***r***, x axis) to rs10783847, consistent with a model of a single causal variant driving both disease association and eQTL.

Among our findings are cases where an eQTL is consistent with associations to multiple diseases. For example, the ankyrin repeat domain 55 (*ANKRD55*) transcript encoded on chromosome 5 has an eQTL in CD4^+^ T cells that is shared with proximal associations to multiple sclerosis, Crohn disease and rheumatoid arthritis (Figure 3, all observations are significant after Bonferroni correction). We also find weaker evidence for shared effects between all three diseases and an eQTL for interleukin 6 signal transducer (*IL6ST*) in CD4^+^ T cells, which passes the false discovery rate threshold but not the more stringent Bonferroni correction (Figure S7). Similarly, a CD4^+^ eQTL for *ELMO1* on chromosome 7 is consistent with associations to both celiac disease and multiple sclerosis (Figure S8), a CD14^+^ eQTL for *RGS1* on chromosome 1 is consistent with associations to both celiac disease and multiple sclerosis (Figure S9), and a CD14^+^ eQTL for *UBE2L3* on chromosome 22 is consistent with associations to both celiac disease and inflammatory bowel disease (Figure S10). In all cases, these are the only genome-wide significant disease associations reported in these loci. As we consider each disease association independently, these results indicate that the same underlying risk variants drive risk to multiple diseases in these loci by altering gene expression, consistent with observations of shared effects across diseases^7^.

**Figure 3.**
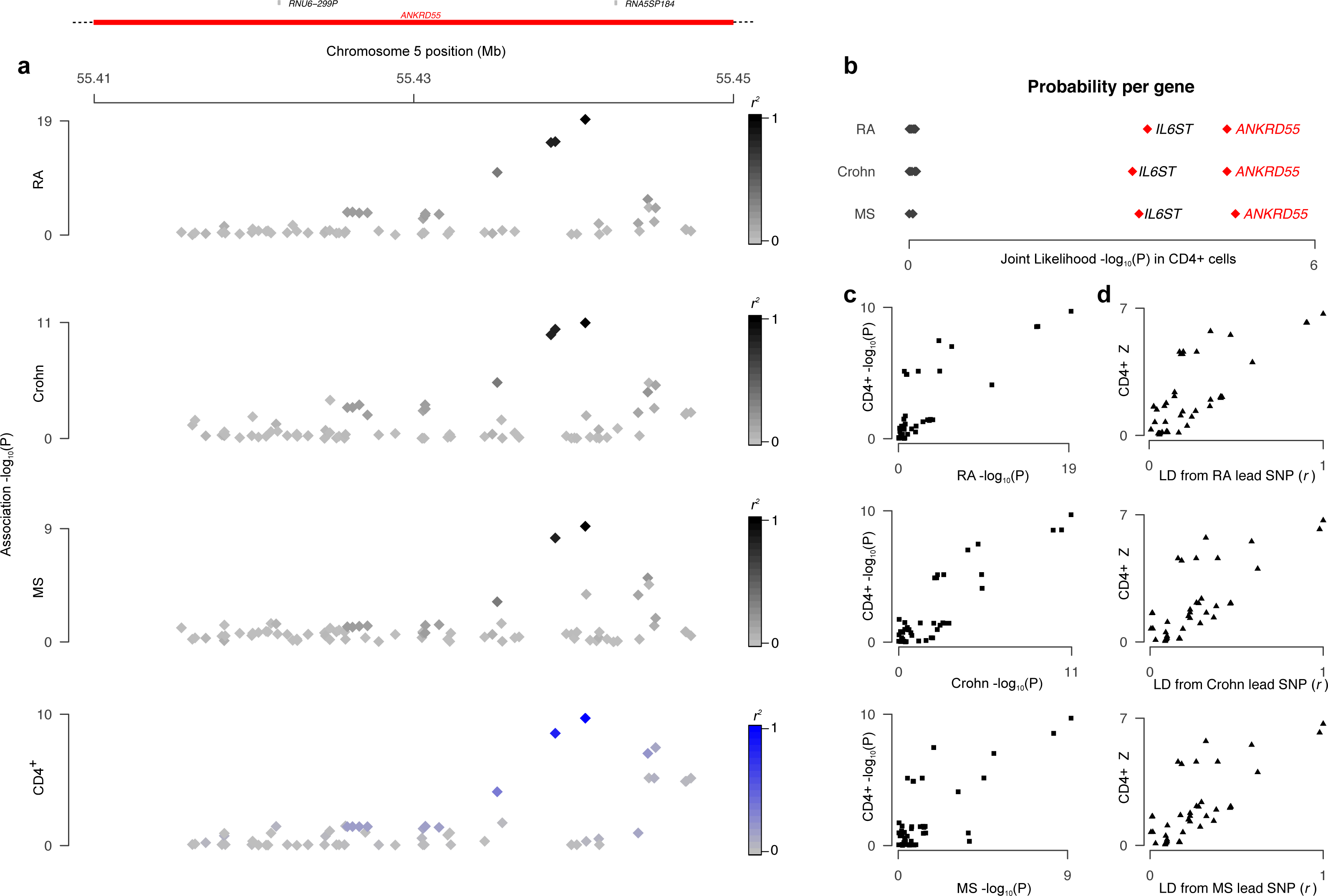
Associations to multiple sclerosis, Crohn disease and rheumatoid arthritis (RA) on chromosome 5 are consistent with an eQTL for *ANKRD55* in CD4 ^+^ T cells. **(a)** Genome-wide significant associations to all three diseases (upper panels) and eQTL data for *ANKRD55* (lower panel; shading in all panels proportional to LD to the most associated variant rs71624119). Due to the variable density of ImmunoChip data, the analysis window is small and only overlaps the coding region of *ANKRD55*, though we test eQTLs for 11 genes with a transcriptional start site within 1Mb of the the association. **(b)** Joint likelihood p-values for nine candidate genes analyzed for this locus in CD4^+^ T cells. Those with FDR < 5% are shown in red. **(c)** Association p-values for each disease (x axis) are strongly correlated to those for the *ANKRD55* eQTL in CD4^+^ cells (y axis). **(d)** Similarly, eQTL association *Z* statistics scale linearly with LD (***r***, x axis) to rs71624119 for all three diseases, consistent with a model of a single causal variant driving all disease associations and the eQTL.

Overall, our results suggest that some autoimmune and inflammatory disease loci are consistent with eQTLs acting in specific immune cell subpopulations, which form strong mechanistic hypotheses for the molecular mechanisms driving disease risk. However, these only account for a small fraction of eQTLs present in disease risk loci; this suggests that abundant caution must be exercised before inferring pathological relevance for an observed eQTL simply due to proximity to a disease association. Strong evidence of a shared genetic effect should therefore be established prior to embarking on time-consuming and costly experimental dissection of such effects.

Previous efforts to detect shared effects between traits in specific loci rely on conditional analyses^29^ or indirectly leverage linkage disequilibrium to test if the shape of association peaks in the region are similar^19,26,27,30^. In contrast, we directly evaluate whether the data support a shared effect through joint likelihood estimation. Through this direct evaluation, we are able to resolve cases where the associations are proximal with higher resolution (Figure S3, Tables S3 and S4). As our method is general, we suggest it may be useful in other contexts, for example in establishing if the shared heritability between diseases is driven by the same underlying causal effects^31^.

More broadly, our results raise the question of how causal disease variants alter cell function to induce risk, given the strong enrichment of disease risk signal on regions of chromatin accessibility with gene regulatory potential^1^, and gene enhancers in particular^14^. We suggest that although gene regulatory regions harboring risk variants are accessible in multiple immune cell subpopulations, they may control gene expression in either a tissue-specific or condition-specific manner, which is not manifest in all cell populations. Our results therefore reinforce the view that we must seek the appropriate cell type and physiological conditions in order to capture the pathologically relevant gene regulatory changes driving disease risk.

## Methods

### Simulated dataset

We selected two loci with previously known associations to base our positive and negative simulations: CD58 on chromosome 1, rs667309 associated to MS risk^32^, and ATG16L1 on chromosome 2, rs2241880 associated to IBD^33^. In each, we used Hapgen^34^ and phased base haplotypes of the locus (2*n*=112, downloaded from http://mathgen.stats.ox.ac.uk/impute/impute_v2.html) to simulate up to ten cohorts, each with 1,000 cases and 1,000 controls, and up to 90 cohorts with other causal variants (Table S1). We chose these other variants to have a range of LD to rs667309 or rs2241880, so we could assess the LD resolution of our joint likelihood method. In each cohort, we simulated genotypes for all variants within 2Mb of rs667309 or rs2241880. To make the cohorts comparable to each other, we adjusted the effect sizes of causal SNPs at different MAF to maintain full power in each simulated cohort. The pairs of simulated cohorts were used either as positive controls if they simulate the same causal variant and negative controls otherwise, to assess the LD resolution limits of our method. Cohort pairs simulating distinct causal variants in high LD (*r^2^* > 0.8) were excluded. For each pair, one cohort was treated as the primary trait (i.e. a disease GWAS) and the other as a secondary trait (i.e. eQTL). Due to the nature of the coalescent forward simulation model, 7.7% of simulated cohorts showed peak association to a SNP in only moderate LD with the specified causal variant (*r^2^* < 0.8). We kept these cohorts as secondary traits only, to better capture the vagaries of resolution limits inherent in the small sample size of eQTL studies. In total, we generated 1,629 cohort pairs for positive controls simulating the same causal variant and 6,106, 530, and 160 cohort pairs for negative controls for which distinct causal variants are separated by *r^2^* of 0 – 0.5, 0.5 – 0.7, and 0.7 – 0.8, respectively.

### Disease GWAS datasets

We downloaded association summary statistics for type 1 diabetes (T1D), rheumatoid arthritis (RA), celiac disease (CEL), multiple sclerosis (MS), inflammatory bowel disease (IBD), Crohn’s disease (Crohn), and ulcerative colitis (UC) from ImmunoBase (immunobase.org; Table S2). For MS, we used the association statistics derived from the combined cohort of discovery and validation samples^8^ in order to maximize the sample size and genetic resolution. For IBD, Crohn, and UC, summary data are from European subset of a trans-ethnic association stud^28^. All association data are solely based on ImmunoChip samples and do not include imputed genotypes. To address population structure, we limited our analyses to European subjects only with the exception of RA^35^, which includes 620 Punjab individuals out of a total of 27,345. T1D summary statistics are from the meta-analysis between case/control association and affected sib-pair analysis.

As our method works best on dense genotype data, we restricted our analyses to the 188 loci genotyped at high density on ImmunoChip. We excluded the Major Histocompatibility Complex (MHC) locus, due to the complex landscape of selection and resulting complex LD patterns. For each disease, we sought the largest published genetic mapping study and identified genome-wide significant associations reported in the 188 ImmunoChip loci. We note that these reports may contain additional samples, so the associations may not be genome-wide significant in the ImmunoChip studies alone. We also excluded any secondary associations after conditioning on initial results, as these are inconsistently reported across diseases. If multiple independent associations are reported within the same ImmunoChip region for any disease, we divide the region at the mid-point between the reported markers and select lead SNPs in each sub-interval separately.

### eQTL dataset

We examined eQTLs in lymphoblastoid cell lines^23^ (LCLs) and primary CD4^+^ T cells and CD14^+^ monocytes^24^ obtained from healthy donors (Table S2). For LCLs, we obtained imputed genotypes and normalized RNAseq in RPKM for 278 non-Finnish European donors in the Geuvadis project. We removed SNPs with minor allele frequency (< 5%), high probability of Hardy-Weinberg disequilibrium (P_HW_ < 10^−5^), or high genotype missing rate (>5%). We removed pseudogenes and transcripts without assigned gene symbols from the expression data, and calculated association statistics by linear regression of genotype on expression levels, including three population principal components to control for structure^36,37^. For CD4^+^ and CD14^+^, we regressed normalized expression levels for European Americans (*n*=213 and 211, respectively) on similarly QCed imputed allele dosages. For all cell types, we generated adaptive permutation statistics from 10^3^ up to 10^6^ iterations^36^, using all covariates.

### Joint likelihood mapping (JLIM)

To test the hypothesis that association signals for two traits are driven by the same causal variant, we contrasted the joint likelihood of observed association statistics under the assumption of same compared to distinct causal variant. Due to limited genetic resolution, distinct causal variants were defined by separation in LD space by *r*^2^ < *θ* from each other. The limit of genetic resolution *θ* is a user-specified parameter and was set to 0.8 in this study. We assumed that at most one causal variant was present in the locus for each trait and that no samples overlap between the traits. We designed the joint likelihood mapping (JLIM) statistic *Λ* in an asymmetrical fashion, requiring only summary-level statistics for one trait (primary trait) but genotype-level data for the other (secondary trait). Specifically, *Λ* was defined as the sum of log likelihood that the causal variant underlying secondary trait is more likely to be same as than distinct from the variant underlying primary trait, as integrated over a set of likely causal variants under a GWAS peak of primary trait:

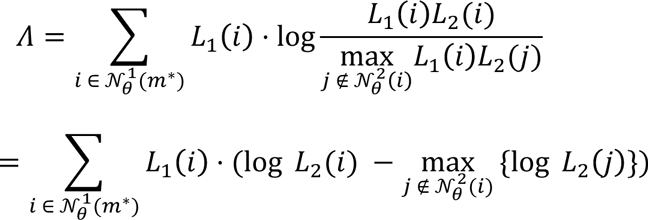

where *m\** is the most associated SNP for primary trait, *L*_1_*(i)* and *L*_2_*(i)* are the likelihood of SNP *i* being causally associated with primary and secondary traits, respectively, and *N_θ_*^1^*(i)* and *N_θ_*^2^*(i)* are the sets of SNPs within LD neighborhood around SNP *i*, as defined by {*SNP_j_|r*_*i*,*j*_^2^ > *θ*}. We derived *N_θ_*^1^ from the reference LD panel and *N_θ_*^2^ directly from the genotypes of secondary trait cohort. We used disease outcome as primary trait, leveraging the larger sample size and dense genotyping, and gene expression as secondary trait, taking advantage of the availability of individual genotype data.

We calculated the likelihood of causal association by approximating the local LD structure with pairwise correlation as previously described^38,39^. Briefly, when SNP *c* is the only causal variant in the locus with non-centrality *λ_c_*, the association static *z_i_* of non-causal SNP *i* follows a normal distribution *N*(*r_i_*_,*c*_ λ_*c*_, 1), where *r_i_*_,*c*_ is LD between SNPs *i* and *c* measured in pairwise Pearson correlation of genotypes. In general, when association statistics ***Z*** *=* (*z*_1_*, z*_2_*,…, z_M_*)*^T^* are provided for all *M* SNPs in the analysis window, the likelihood of SNP *i* being the causal variant with non-centrality *λ_i_* is:

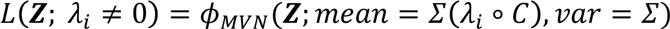

where *ϕ_MVN_* is the multivariate normal density function, *C* is an incident vector with *C_k_ = 1* if and only if *k = i*, *Σ* is a *M* x *M* local LD matrix defined by pairwise Pearson correlation between genotypes, and ο is element-wise multiplication [Kichaev]. Since we do not know the true non-centrality of causal variant, we estimated the profile likelihood, which simplifies to a closed form:

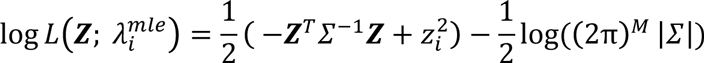

with *λ_i_^mle^ = z_i_*. Thus, given association statistics for primary and secondary traits, ***Z*** = (*z*_1_*, z*_2_*,…, z_m_*)*^T^* and ***W*** *=* (*w*_1_*, w*_2_*,…, w_m_*)*^T^*, the test statistic *Λ* simplifies to:

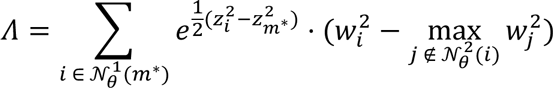

The p-value of joint likelihood is estimated by permuting phenotypes of secondary traits as under the trivial null hypothesis that that there is no casual variant for secondary trait in the locus (“*H*_0_”). With respect to the more likely null that distinct causal variants underlie association signals of two traits (“*H*_2_”), we can show that asymptotically as the non-centrality of causal variant increases, p-values estimated from *H*_0_ behave conservatively with respect to *H*_2_ (Supplementary Notes):

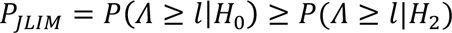

Thus, with large enough sample or effect sizes, joint likelihood test against *H*_0_ will also reject *H*_2_ in favor of alternative hypothesis of shared causal variant (“*H*_1_”). Further, to evaluate whether this property holds for practical non-centrality values, we examined the negative controls simulating *H*_2_ in ATG16L1 and CD58 loci, specifically, if *P_JLIM_* was highly shifted toward 1.0 (Figure S2) and larger than empirically estimated false positive rates as expected (Table S4).

For both simulated and real GWAS data, we applied JLIM to SNPs with data for both primary and secondary traits, present in the reference LD panels, and within 100kb of the most associated marker to disease (“lead SNP”). In ImmunoChip data, the analysis windows were further confined by the boundaries of the fine-mapping intervals. We compared each lead SNP to eQTL data for all genes with a transcription start sites (TSS) up to 1Mb from the lead SNP, and an eQTL association *p < 0.05* for at least one SNP in the analysis window. To minimize computational burden, we did not consider SNPs associated with neither disease or eQTL (association p > 0.1 to both). For the reference LD panel, we used the base haplotypes of Hapgen simulation for simulated datasets, and non-Finnish European samples (*n*=404) of the 1000 Genomes Project (phase 3, release 2013/05/02) for ImmunoChip loci.

We corrected for multiple tests using false discovery rate (FDR) levels and Bonferroni correction. The FDR was calculated separately for specific disease and cell type combination as:

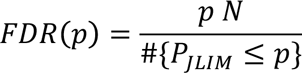

where *p* is a JLIM p-value cut-off, and *N* is the number of all tested disease lead SNP-eQTL candidate gene combinations. The FDR was calculated for each cell type since the distribution of JLIM p-values can vary depending on the disease relevance of cell type. To provide a list of higher confidence hits in each disease, we also applied the Bonferroni correction to nominal JLIM p-values for the number of tests across all three cell types.

### Bayesian *coloc*

We ran Bayesian *coloc*^19^ using default parameter settings on both simulated and real data as described for JLIM. For simulated data, we used colocalization prior *p_12_* values of 10^−5^ or 10^−6^, which are default values for higher sensitivity and higher specificity, respectively. The beta and variance of beta were used for all SNPs in the analysis window in case/control mode. We calculated accuracy as the area under the receiver operator curve (ROC; Figure S3).

As ImmunoChip data is only available as summary statistics, we used the minor allele frequencies from non-Finnish Europeans from the 1000 Genomes Project, and quantitative beta and variance of beta calculated on eQTL association data, and a colocalization prior *p_12_* = 10^−6^. We did not consider the type 1 diabetes data, where case/control sample size is limited after excluding affected sib pair data.

### Estimating the number of disease GWAS loci with consistent eQTL effects

We expect JLIM p-values to follow a bimodal distribution with modes close to zero and one when the data support a model of shared or distinct causal effects, respectively. Conversely, under the null model of no *cis*-eQTL association, we expect a uniform p-value distribution. We can thus estimate the proportion disease-eQTL pairs belonging to the null *π*_0_, same *π*_1_ and distinct *π*_2_ causal variant models from the observed p-value distribution^40^ (Figures S11-13). To assess if the strength of the eQTL association influences the likelihood of identifying a shared causal variant, we calculate these proportions for subsets of trait pairs defined by minimum eQTL p-value. In each bin, we identified the limits of the uniform portion of the distribution *γ*_1_ and *γ*_2_ and estimate *π*_0_, *π*_1_, and *π*_2_ as:

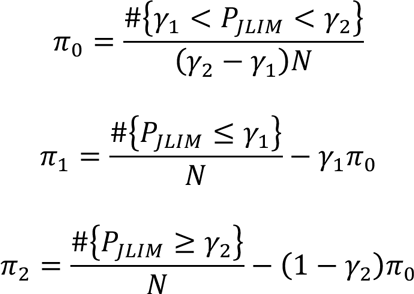

To estimate the number of disease GWAS loci that can be explained by consistent effect of same causal variant on disease and eQTL (denoted by *C* below), we incrementally relaxed the p-value cut-offs of JLIM and examined the trends of the number of disease loci with at least one JLIM hit and subtracted the expected number of false positive loci (Figures 1 and S14). Specifically, at each JLIM p-value cutoff *p*_!_, we successively calculated *C*(*p_i_*):

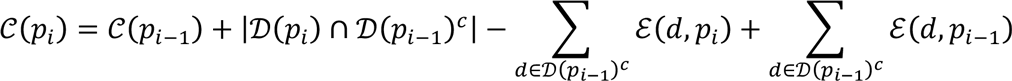

where *p_i_*_−1_ < *p*_i_ with *p*_0_ = 0, *D*(*p*) is the set of disease GWAS loci with at least one eQTL gene in any cell type passing the JLIM p-value cut-off *p*, and &#x2130;(*d*, *p*) is the probability that disease GWAS locus *d* has a false positive eQTL gene passing the JLIM p-value cutoff *p*. We estimated the lower and upper bounds of &#x2130;(*d*, *p*) using the Monte Carlo method by randomly selecting false positive eQTL genes within the locus *d* at rates of (*1 – π*_1_)· *lb* or (*1 – π*_1_) · *ub* over 1,000 iterations. The *lb* and *ub* are the lower and upper bounds of false positive rate of JLIM against true null. Note that *π*_1_ and *lb* depend on the cell type and strength of eQTL association.

As the true null is mixture of two nulls, *H*_0_ and *H*_2_, the false positive rate of JLIM against true null *P* (*Λ ≥ l* |*H*_0_ ∪ *H*_2_) can be bounded by using the following decomposition:

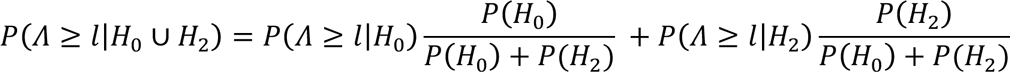

While the false positive rate under distinct null *P* (*Λ ≥ l* |*H*_2_) is difficult to estimate, it is non-negative by definition and asymptotically bounded by permutation p-value *P* (*Λ ≥ l* |*H*_0_), i.e. *P_JLIM_*, as the non-centrality of causal variant increases. Therefore, we took:

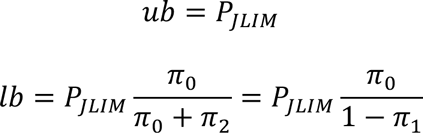

and estimated the bounds of locus-level false positive rates &#x2130;(*d, p*) and number of disease loci with consistent effects *C*(*p_i_*).

